# Using short-read 16S rRNA sequencing of multiple variable regions to generate high-quality results to a species level

**DOI:** 10.1101/2024.05.13.591068

**Authors:** Amy S Graham, Fadheela Patel, Francesca Little, Andre van der Kouwe, Mamadou Kaba, Martha J Holmes

## Abstract

**Introduction:** Short-read amplicon sequencing studies have typically focused on 1-2 variable regions of the 16S rRNA gene. Species-level resolution is limited in these studies, as each variable region enables the characterisation of a different subsection of the microbiome. Although long-read sequencing techniques take advantage of all 9 variable regions by sequencing the entire 16S rRNA gene, they are substantially more expensive. This work assessed the feasibility of accurate species-level resolution and reproducibility using a relatively new sequencing kit and bioinformatics pipeline developed for short-read sequencing of multiple variable regions of the 16S rRNA gene. In addition, we evaluated the potential impact of different sample collection methods on our outcomes.

**Methods:** Using xGen^TM^ 16S Amplicon Panel v2 kits, sequencing of all 9 variable regions of the 16S rRNA gene was carried out on an Illumina MiSeq platform. Mock cells and mock DNA for 8 bacterial species were included as extraction and sequencing controls respectively. Within-run and between-run replicate samples, and pairs of stool and rectal swabs collected at 0-5 weeks from the same participants, were incorporated. Observed relative abundances of each species were compared to theoretical abundances provided by ZymoBIOMICS. Paired Wilcoxon rank sum tests and distance-based intraclass correlation coefficients were used to statistically compare alpha and beta diversity measures, respectively, for pairs of replicates and stool/rectal swab sample pairs.

**Results:** Using multiple variable regions of the 16S ribosomal Ribonucleic Acid (rRNA) gene, we found that we could accurately identify taxa to a species level and obtain highly reproducible results at a species level. Yet, the microbial profiles of stool and rectal swab sample pairs differed substantially despite being collected concurrently from the same infants.

**Conclusion:** This protocol provides an effective means for studying infant gut microbial samples at a species level. However, sample collection approaches need to be accounted for in any downstream analysis.

## 1. Introduction

Our ability to describe and appreciate the complexities of the human microbiome has been radically improved by next generation sequencing tools (Bharti and Grimm, 2021, Ji and Nielsen, 2015, Rogers and Bruce, 2010). Although the use of second-generation, short-read sequencing platforms allows high read depths to be rapidly sequenced (Hu et al., 2021, Tucker et al., 2009), it is limited in terms of the assembly of contiguous sequences (Li et al., 2010, Zerbino and Birney, 2008). Various third-generation sequencing techniques exist and allow long reads to be sequenced. However, these techniques can be expensive and/or result in higher error rates (Amarasinghe et al., 2020, Midha et al., 2019, Van Dijk et al., 2018). As a result, there is a need to consider alternative ways to improve short-read sequencing approaches.

The 16S rRNA gene has been identified as a particularly useful target of research as it is common to all bacteria (Acinas et al., 2004, Patel, 2001). The gene consists of regions of DNA in which the sequence is conserved across all bacteria, while in other regions there is variation according to the individual bacterial species (Wang and Qian, 2009, Lane et al., 1985). As such, targeted amplicon sequencing can be done, comparing variable region sequences to a database of known taxa, to identify which bacterial species are present in a sample (Wang and Qian, 2009). In the past, amplicon sequencing studies have typically focused on one or two variable regions at a time (Claassen-Weitz et al., 2018, Gao et al., 2018, Yu et al., 2017, Hosgood III et al., 2014, Caporaso et al., 2011, Zhou et al., 2011). Yet, certain variable regions are better for enabling classification to lower taxonomic levels and each variable region favours classification of specific taxa (Bukin et al., 2019, Guo et al., 2013, Chakravorty et al., 2007). Consequently, this approach limits the ability to obtain accurate species-level resolution when focusing only on a single short fragment of the 16S rRNA gene. Using the entire 16S rRNA sequence is expected to provide better classification potential to a species level (Johnson et al., 2019).

The use of short-read sequencing techniques to study multiple variable regions of the 16S rRNA gene has captured the interest of researchers. There has been a rapid development in sequencing kits and bioinformatics pipelines to process multiple variable region 16S rRNA sequencing data (Callahan et al., 2021, Fuks et al., 2018, Schriefer et al., 2018, Wang et al., 2016, Amir et al., 2013). The xGen^TM^ 16S Amplicon Panel v2 kits (Integrated DNA Technologies, Coralville, IA, USA) are an example, having been developed to amplify all nine variable regions of the 16S rRNA gene. Furthermore, a complementary bioinformatics pipeline known as the Swift Normalase Amplicon Panels APP for Python 3 (SNAPP-py3), was developed specifically for the analysis of sequencing data obtained using these kits (Chai, 2021).

Being relatively new, there are only a few publications in which the SNAPP-py3 pipeline has been used to analyse data sequenced with the xGen kits (Nuccio et al., 2023, Bennato et al., 2022). However, neither of these studies took advantage of the species-level classification that can be achieved with the xGen kits and SNAPP-py3 pipeline. Although Bennato et al. (2022) included a control containing DNA for 20 known bacterial species, they only reported the ability to pick up these bacteria at a genus level. To our knowledge, the combined ability of these kits and pipeline to obtain accurate species-level classification has not been assessed. Therefore, we sought to establish a protocol in which the SNAPP-py3 pipeline and additional processing steps were utilised to analyse short-read multiple variable region 16S rRNA data following sequencing with xGen amplicon panel kits.

The accuracy of sequencing protocols can be evaluated in a few ways using mock controls. Firstly, researchers can calculate the proportion of expected species that have been detected down to a species level when using a given protocol and for select regions of the 16S rRNA gene (Johnson et al., 2019, Fouhy et al., 2016). F-scores can be calculated based on the precision and sensitivity with which these species are identified (Özkurt et al., 2022). The classification process can also be assessed according to the percentage of overall reads that are classified as belonging to one of the expected control species (Szoboszlay et al., 2023, Urban et al., 2021). Furthermore, accuracy can be assessed by comparing observed relative abundances to expected abundances (provided by suppliers) for each taxon in a control (Maki et al., 2023, Szoboszlay et al., 2023, Drengenes et al., 2021, Laursen et al., 2017, Caporaso et al., 2011). This can be done at different taxonomic levels and gives an indication of whether the amplification or sequencing processes have introduced bias by favouring certain species over others.

As stool collection is not always possible due to various factors, rectal swab collection has become a common sampling method for studying the gut microbiome (Bassis et al., 2017). Storage of rectal swabs differs to that of stool, as swabs generally need to be placed in a medium (CDC, 2015), for example PrimeStore (Flygel et al., 2020). The results obtained from sequencing rectal swab samples can be inconsistent in terms of numbers of bacteria detected (Chanderraj et al., 2022).

Previous studies have explored whether rectal swab samples can provide a reliable alternative to stool samples (Radhakrishnan et al., 2023, Bokulich et al., 2019, Reyman et al., 2019, Bassis et al., 2017, Freedman et al., 2017). Although pairs of stool and rectal swab samples collected concurrently from the same individual generally display similar diversity and functional profiles (Radhakrishnan et al., 2023, Reyman et al., 2019, Bassis et al., 2017), there have been other studies that suggest that these samples are not equivalent for detecting specific taxa (Jones et al., 2018, Freedman et al., 2017, Goldfarb et al., 2014). In particular, rectal swab samples have been found to be more effective for detecting a greater number of harmful species in children with gastrointestinal infections (Freedman et al., 2017, Goldfarb et al., 2014). When sequencing the meconium (first stool) sample passed by newborn infants, rectal swab samples have been found to provide a less accurate representation of the microbiome compared to stool samples (Graspeuntner et al., 2023). Moreover, rectal swab samples provide a poorer representation of the microbiome if they are sequenced after greater than 48 hours at room temperature (Bokulich et al., 2019). As a result, the interchangeability of stool and rectal swab samples needs to be assessed for new protocols. Moreover, sample collection approach is an important variable to consider with regards to the research objectives of a study.

The first aim of this research involves assessing the accuracy of extraction and sequencing protocols to achieve classification at the species level. This will be done by sequencing and analysing mock controls containing either whole cells or already-extracted DNA from eight known bacterial species, assessing the precision and sensitivity with which species were identified and how well their relative abundances matched theoretical abundances. Secondly, our goal is to evaluate the within-run and between-run reproducibility of species-level analysis. Technical replicate samples sequenced on either the same plate (within-run) or across different sequencing plates and runs (between-run), will be compared to determine this. Finally, we aim to identify whether there are differences at a species level between different sample collection approaches. To achieve this, we will compare pairs of stool and rectal swab samples collected from the same participants at the same time point (0-5 week-old newborns). We hypothesise that by sequencing multiple variable regions of the 16S rRNA gene and using the SNAPP-py3 pipeline, we could obtain accurate species-level resolution and achieve reproducible results.

## 2. Results

Analysis of mock controls and technical replicates was carried out to assess the use of the xGen Amplicon kits and the SNAPP-py3 pipeline as a multivariate 16S rRNA sequencing approach for achieving accurate and reproducible species-level resolution. Furthermore, the similarity of samples collected using different sample collection techniques was investigated by comparing pairs of baseline stool and rectal swab samples from the same participants.

### 2.1 DNA extraction reliability

All eight expected bacterial species were detected in four of our seven mock extraction controls (Table 1). *Bacillus subtilis* was not detected to the species level in two controls (Table 2; Figure 1a), however classification to a genus level (*Bacillus*) was achieved (Supplementary table 1). *Listeria monocytogenes* was not detected even at a genus level in the control from run 2, plate 1 (Supplementary table 1). For the three controls in which we were unable to detect all eight species, the total percentage of sequencing data correctly classified as expected mock species was consequently lower (Table 2). Sensitivity scores were over 0.88 for all controls. Precision scores were lower – particularly for the control on run 1, plate 1, which had a score of 0.32. A median F-score of 0.84 (range of 0.47 to 1.00) was obtained for the mock extraction controls (Table 1).

**Figure 1:**
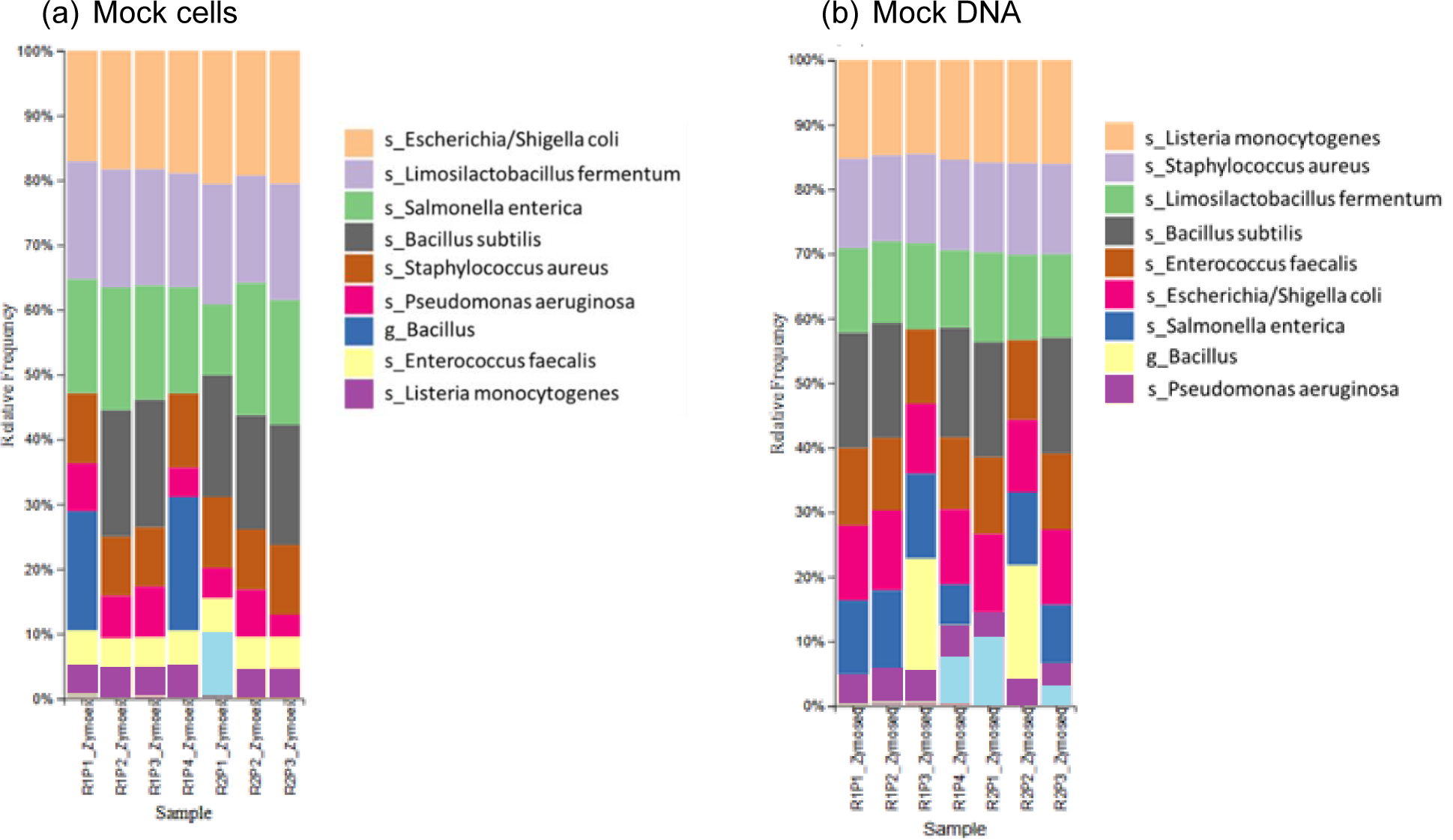
Relative abundances for (a) mock cell and (b) mock DNA controls; Legend: s = species, g = genus level classification. R# = run number; P# = plate number. The legend only includes taxa of interest to the species or genus level.

**Table 1:** A summary of performance and accuracy measures at a species level for mock cell controls containing eight known bacterial species. R# = run number; P# = plate number.

|  | R1P1<br>Zymoex | R1P2<br>Zymoex | R1P3<br>Zymoex | R1P4<br>Zymoex | R2P1<br>Zymoex | R2P2<br>Zymoex | R2P3<br>Zymoex |
| --- | --- | --- | --- | --- | --- | --- | --- |
| True positives | 7 | 8 | 8 | 7 | 7 | 8 | 8 |
| Precision | 0.32 | 0.73 | 0.73 | 1.00 | 0.70 | 1.00 | 0.80 |
| Sensitivity | 0.88 | 1.00 | 1.00 | 0.88 | 0.88 | 1.00 | 1.00 |
| F-score | 0.47 | 0.84 | 0.84 | 0.93 | 0.78 | 1.00 | 0.89 |

**Table 2:**
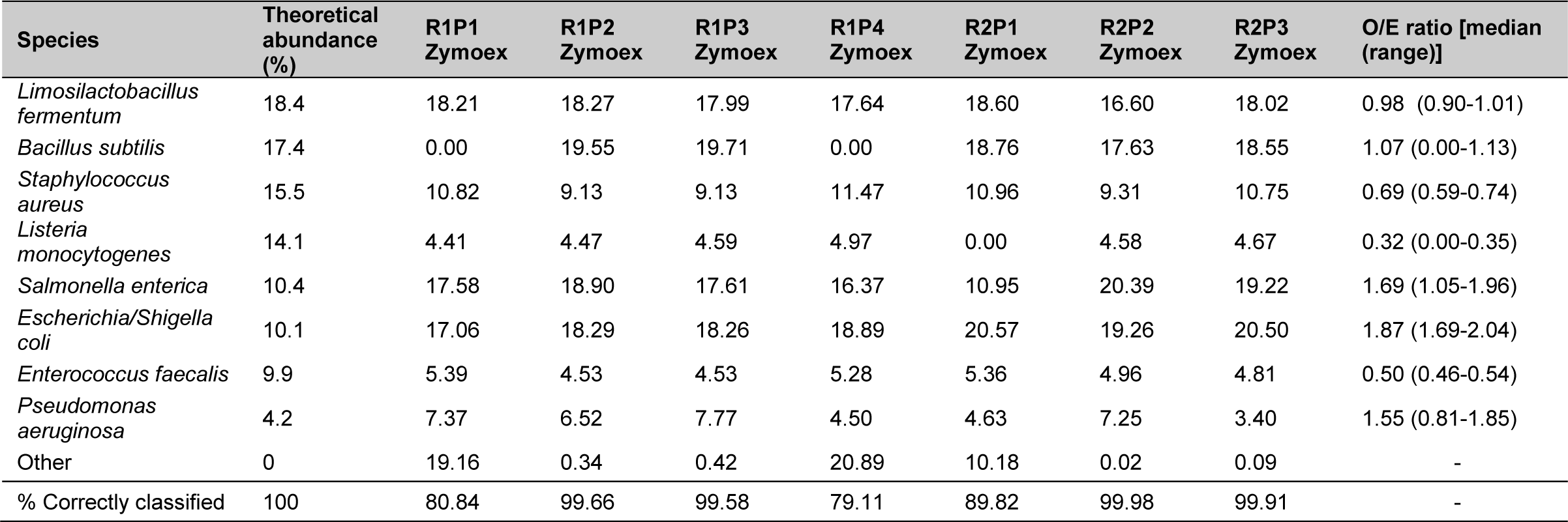
The theoretical abundances and relative abundances of the eight expected bacterial species in mock cell controls given as percentages. The median observed/expected (O/E) ratios are also provided. The total percentage of sequences correctly classified as one of the expected species is given for each control. R# = run number; P# = plate number.

The percentage abundances of species in these controls did not accurately follow the order of theoretical abundances for some species (Table 2; Figure 2a). In particular, the relative abundances of *Listeria monocytogenes* were well below the theoretical abundances suggested by ZymoBIOMICS at both a species and genus level. This is emphasised by the low median Observed/Expected (O/E) ratio of 0.32 at a species level. Similarly, *Enterococcus faecalis* and *Staphylococcus aureus* had O/E ratios well below the value of 1. The relative abundances of *Escherichia coli* and *Salmonella enterica* were greater than expected, with a range of O/E ratios all lying well above the value of 1.

**Figure 2:**
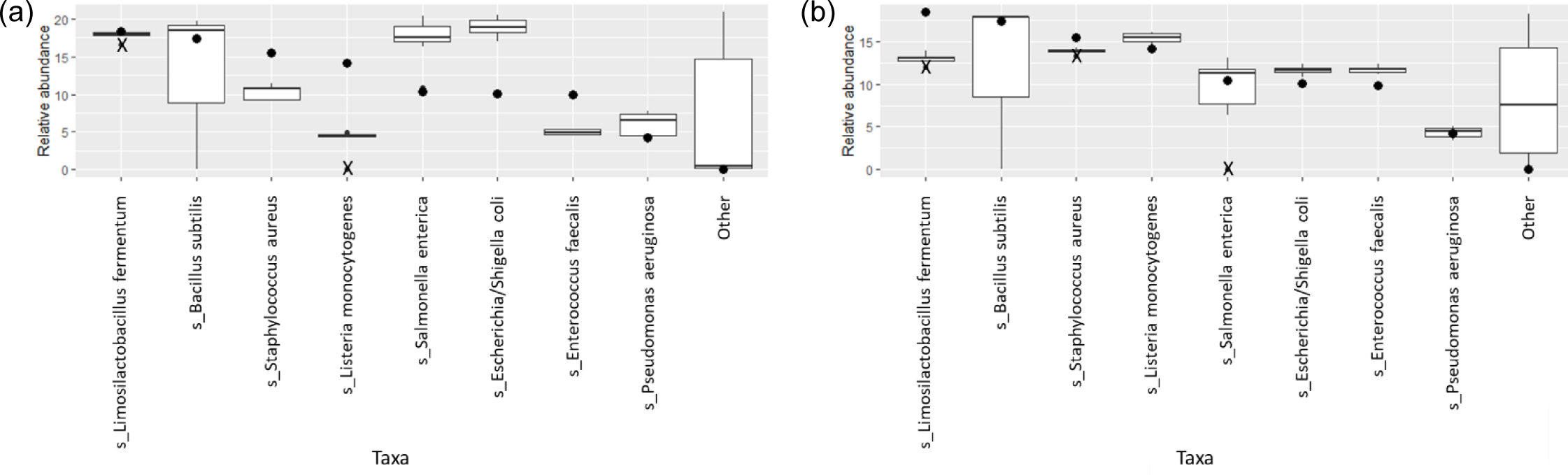
Box and whisker plots showing the relative abundance percentages of the eight expected bacterial species across (a) mock extraction controls and (b) mock DNA controls; theoretical abundances shown as dots; outliers shown as crosses.

### 2.2 Sequencing reliability

Among the mock sequencing controls, the eight anticipated bacterial species were detected in five of the seven controls (Table 3; Figure 1b). The overall percentages of sequences correctly classified as expected species were slightly lower for these controls compared to the mock extraction controls (Table 4). For the run 2 plate 1 control the prevalence of *Salmonella enterica* was particularly low, and this was not resolved at the genus level (Supplementary table 2). *Bacillus subtilis* again was not detected at a species level for two of these controls (Table 3). Precision scores for the mock sequencing controls ranged from 0.50 and up, while sensitivity was greater than 0.88 for all controls. F-scores had a median of 0.80 (range of 0.67 to 0.94).

**Table 3:** A summary of performance and accuracy measures at a species level for mock DNA controls containing eight known bacterial species. R# = run number; P# = plate number.

|  | R1P1<br>Zymoseq | R1P2<br>Zymoseq | R1P3<br>Zymoseq | R1P4<br>Zymoseq | R2P1<br>Zymoseq | R2P2<br>Zymoseq | R2P3<br>Zymoseq |
| --- | --- | --- | --- | --- | --- | --- | --- |
| True positives | 8 | 8 | 7 | 8 | 8 | 7 | 8 |
| Precision | 0.67 | 0.50 | 0.64 | 0.57 | 0.89 | 1.00 | 0.73 |
| Sensitivity | 1.00 | 1.00 | 0.88 | 1.00 | 1.00 | 0.88 | 1.00 |
| F-score | 0.80 | 0.67 | 0.74 | 0.73 | 0.94 | 0.93 | 0.84 |

**Table 4:** The theoretical abundances and relative abundances of the eight expected bacterial species in mock DNA controls are given as percentages. The median observed/expected (O/E) ratios are also provided. The total percentage of sequences correctly classified as one of the expected species is given for each control. R# = run number; P# = plate number.

| Species | Theoretical abundance (%) | R1P1 Zymoseq | R1P2 Zymoseq | R1P3 Zymoseq | R1P4 Zymoseq | R2P1 Zymoseq | R2P2 Zymoseq | R2P3 Zymoseq | O/E ratio [median (range)] |
| --- | --- | --- | --- | --- | --- | --- | --- | --- | --- |
| *Limosilactobacillus fermentum* | 18.4 | 13.08 | 12.60 | 13.32 | 11.94 | 13.88 | 13.16 | 12.86 | 0.71 (0.65-0.75) |
| *Bacillus subtilis* | 17.4 | 17.89 | 17.84 | 0.00 | 17.05 | 17.81 | 0.00 | 17.99 | 1.02 (0.00-1.03) |
| *Staphylococcus aureus* | 15.5 | 13.84 | 13.34 | 13.83 | 14.04 | 13.95 | 14.22 | 14.03 | 0.90 (0.86-0.92) |
| *Listeria monocytogenes* | 14.1 | 15.27 | 14.76 | 14.54 | 15.41 | 15.85 | 15.95 | 16.05 | 1.09 (1.03-1.14) |
| *Salmonella enterica* | 10.4 | 11.49 | 12.04 | 13.12 | 6.32 | 0.01 | 11.23 | 9.06 | 1.08 (0.00-1.26) |
| *Escherichia/Shigella coli* | 10.1 | 11.59 | 12.40 | 10.84 | 11.62 | 12.18 | 11.20 | 11.64 | 1.15 (1.07-1.23) |
| *Enterococcus faecalis* | 9.9 | 11.98 | 11.18 | 11.50 | 11.12 | 11.89 | 12.40 | 11.76 | 1.19 (1.12-1.25) |
| *Pseudomonas aeruginosa* | 4.2 | 4.44 | 5.10 | 4.64 | 4.94 | 3.72 | 4.09 | 3.39 | 1.06 (0.81-1.21) |
| Other | 0 | 0.42 | 0.74 | 18.20 | 7.54 | 10.72 | 17.74 | 3.21 | - |
| % Correctly classified |  | 99.58 | 99.26 | 81.80 | 92.46 | 89.28 | 82.26 | 96.79 | - |

The relative abundances of species in the mock sequencing controls more closely matched the expected abundances than was observed for the mock extraction controls (Table 4, Figure 2b), as seen by the O/E ratios being closer to 1. In these controls the abundance of *Limosilactobacillus fermentum* was well below the theoretical threshold expected. The O/E ratios for this species and *Staphylococcus aureus* were consistently less than 1. Whereas *Enterococcus faecalis*, *Escherichia coli* and *Listeria monocytogenes* had O/E ratio ranges above 1.

Although the inclusion of a batch effect correction step was trialled, substantial differences in the relative abundances of the various species across the controls were observed compared to when no batch effect correction was done (Supplementary figure 1). Similarly, stricter decontamination thresholds led to poorer reproducibility in mock controls.

### 2.3 Within-run and between-run reproducibility

Similar patterns in the relative abundance of species could be seen when comparing pairs of within-run and between-run repeats (Supplementary figures 2 & 3). When comparing alpha diversity of within-run repeats using paired Wilcoxon tests, we found no evidence to indicate differences at a species level in the Observed richness (95% confidence interval (CI) [-3.50, 5.50]; p=0.587), Shannon’s index (95% CI [-0.09, 0.12]; p=0.782) or Simpson’s index (95% CI [-0.02, 0.01]; p=0.487) between these pairs of technical replicates (Figure 3). Furthermore, when comparing the beta diversity distance matrices of these within-run technical replicate pairs (Figure 4), we observed a good level of reproducibility for Bray Curtis as seen by a distance-based intraclass correlation coefficient (dICC) value of 0.940 and Jaccard’s distance showed good, albeit lower, reproducibility with a dICC of 0.762. Similarly, we found no differences between these technical replicates when looking at their alpha and beta diversity measures at a genus level.

**Figure 3:**
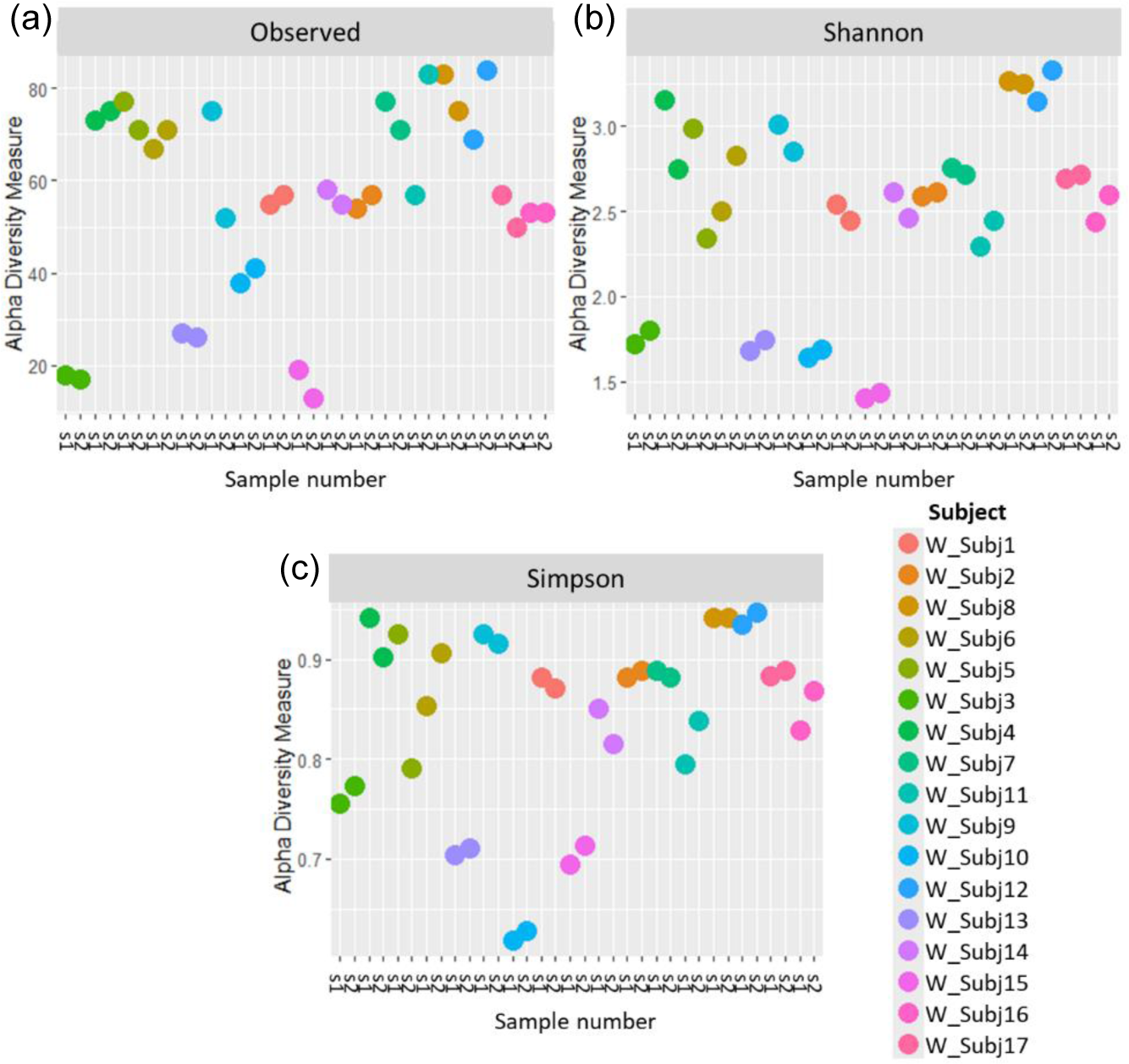
(a) Observed richness, (b) Shannon’s index and (c) Simpson’s index for within-run technical replicates at species-level classification. s1 = sample 1; s2 = sample 2.

**Figure 4:**
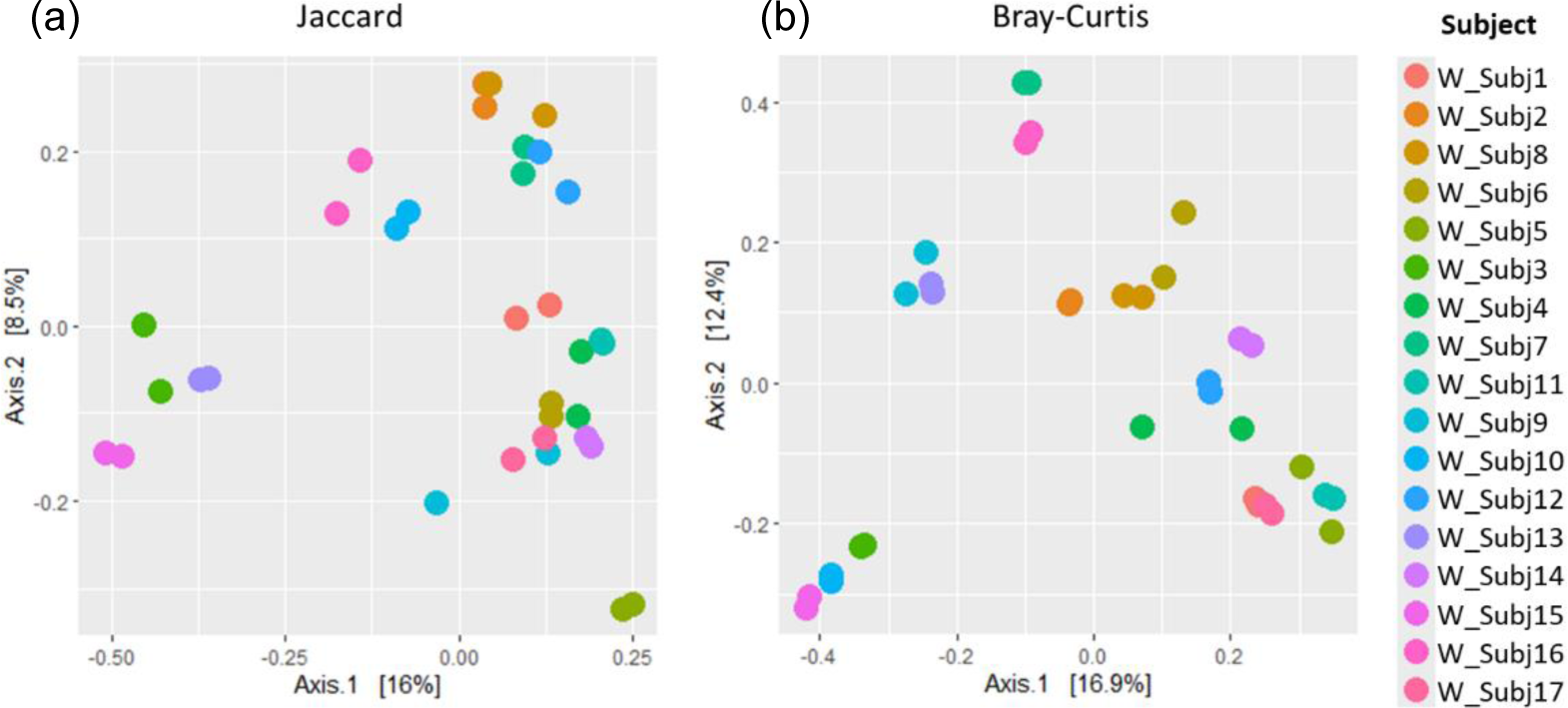
A comparison of (a) Jaccard and (b) Bray Curtis beta diversity measures between pairs of within-run technical replicates at species-level classification.

A comparison of alpha and beta diversity measures for between-run technical replicates similarly found no clear differences between these pairs at a species level (Figure 5). Paired Wilcoxon tests comparing Observed species (95% CI [-2.00, 13.00]; p=0.362), Shannon’s index (95% CI [-0.05, 0.49]; p=0.195) and Simpson’s index (95% CI [-0.004, 0.11]; p=0.148) did not motivate for the existence of differences in alpha diversity measures. Moreover, dICC results showed good reproducibility for Bray Curtis (dICC=0.899) and moderate reproducibility for Jaccard’s distance (dICC=0.597) (Figure 6).

**Figure 5:**
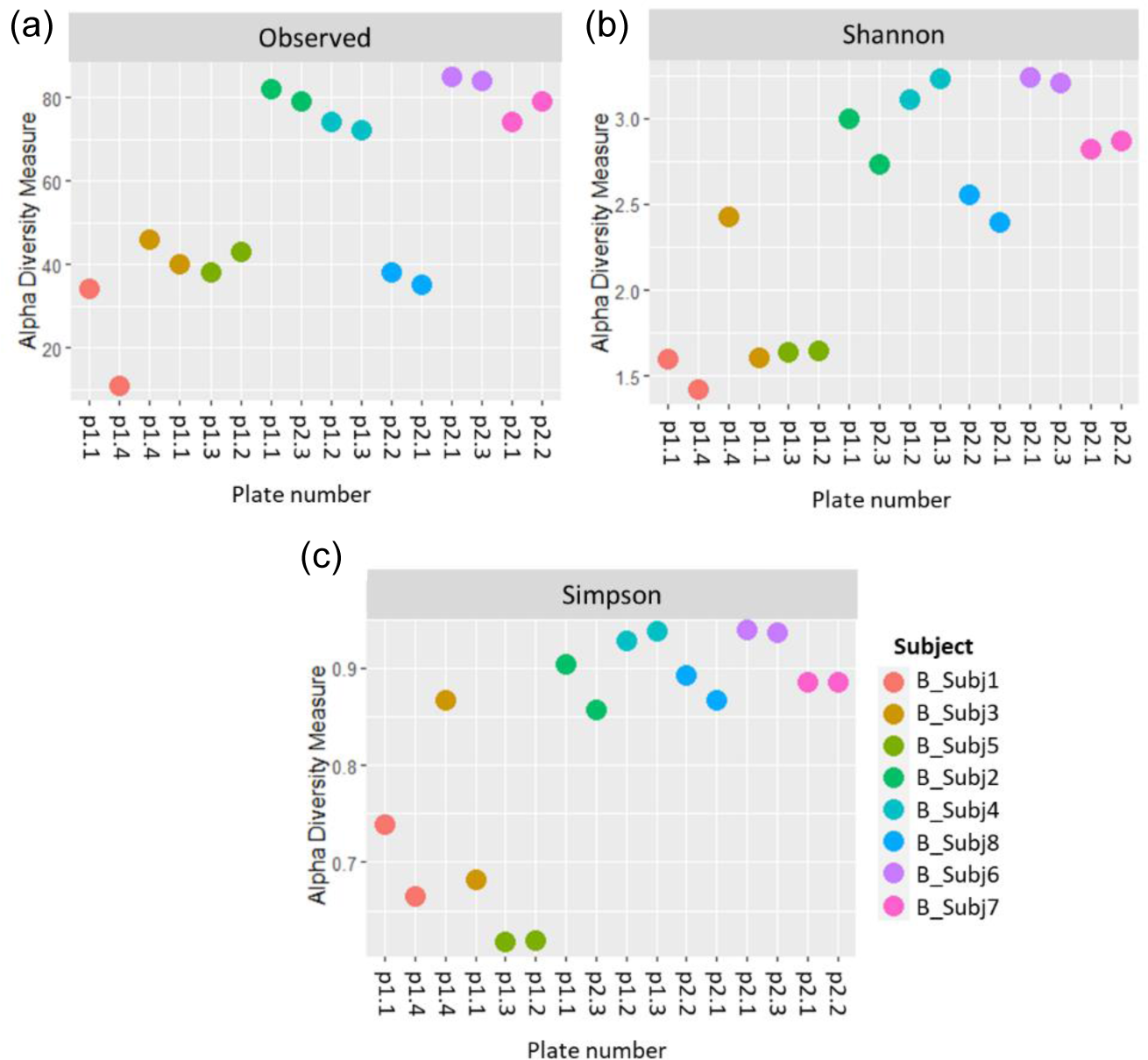
(a) Observed richness, (b) Shannon’s index and (c) Simpson’s index for between-run technical replicates at species-level classification. p# = run & plate number

**Figure 6:**
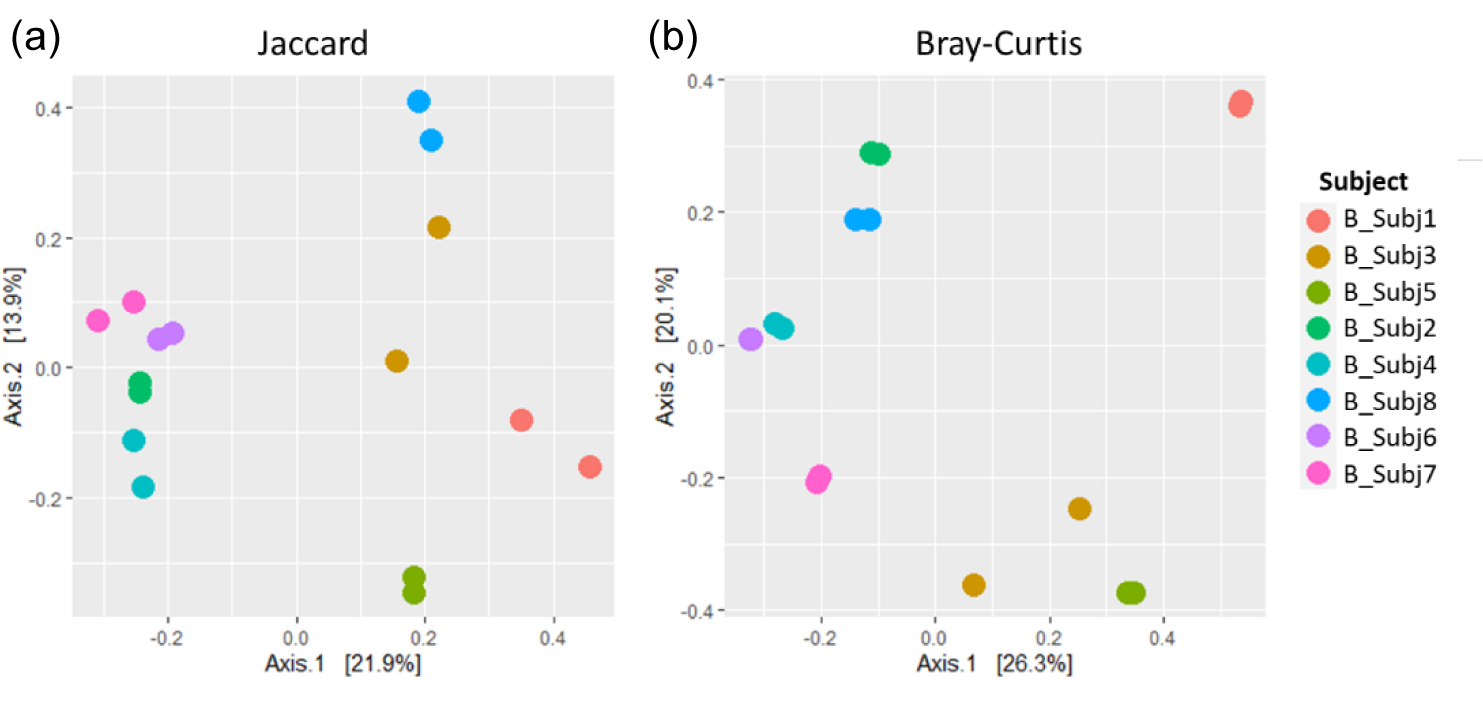
A comparison of (a) Jaccard and (b) Bray Curtis beta diversity measures between pairs of between-run technical replicates at species-level classification.

### 2.4 Interchangeability of stool and rectal swab samples

To assess whether stool and rectal swab samples may be combined for analysis, we compared 26 sample pairs collected at baseline (0 - 5 weeks after birth). The relative abundances of species did not display similar patterns across pairs of samples (Supplementary figure 4) and alpha diversity was found to differ when running paired Wilcoxon tests. Stool and swab samples from the same participants differed at a species level in terms of Observed species (p<0.001) and Shannon’s index (p= 0.027), with no differences in Simpson’s index (p= 0.394) found (Figure 7). A comparison of beta diversity measures at a species level found that while Bray Curtis measures were moderately reliable between the pairs of stool and swab samples (dICC=0.684), there was poor reliability when comparing their Jaccard’s distances (dICC=0.310) (Figure 8).

**Figure 7:**
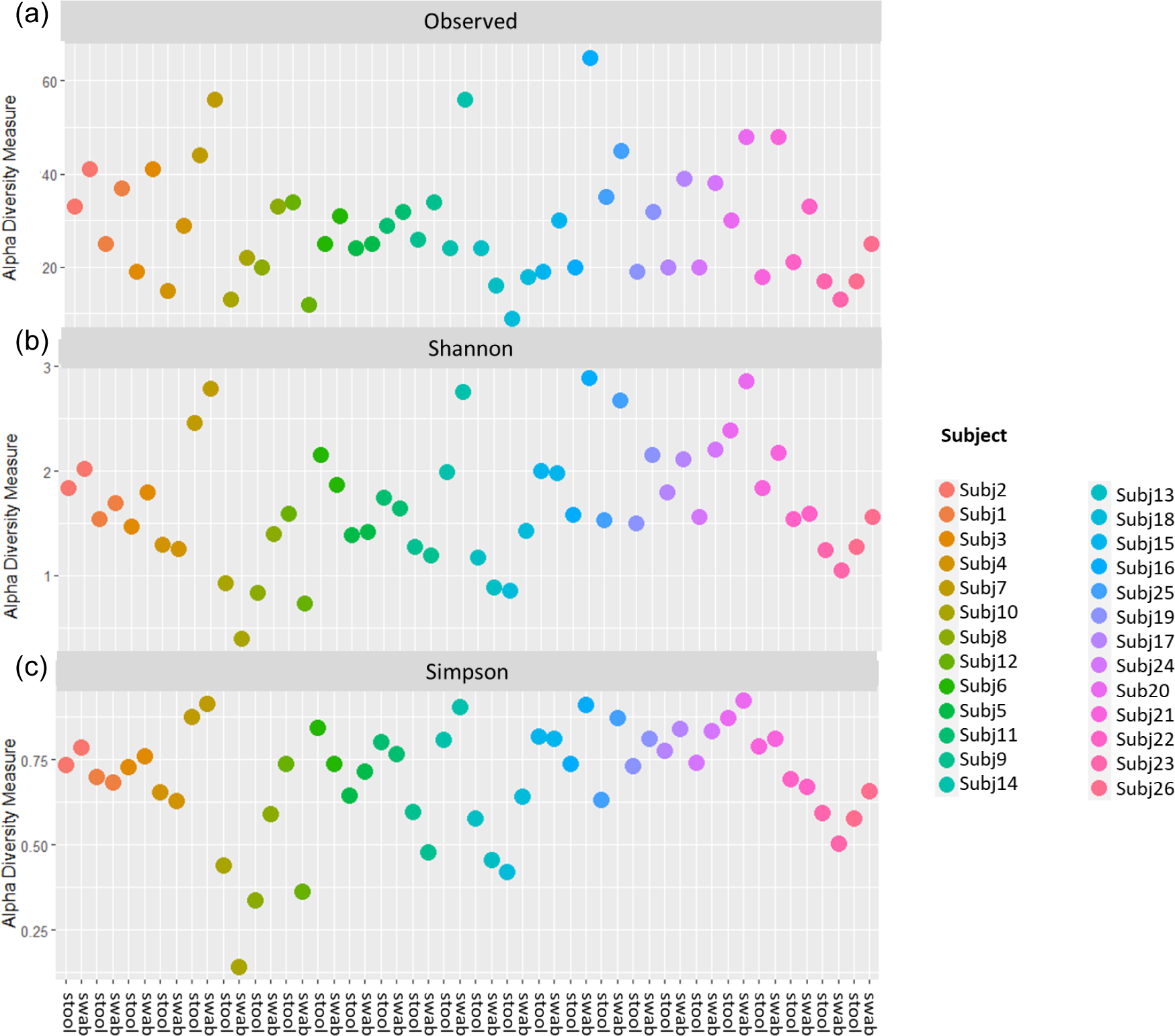
(a) Observed richness, (b) Shannon’s index and (c) Simpson’s index for pairs of stool and rectal swab samples collected from the same infants using data classified to a species level.

**Figure 8:**
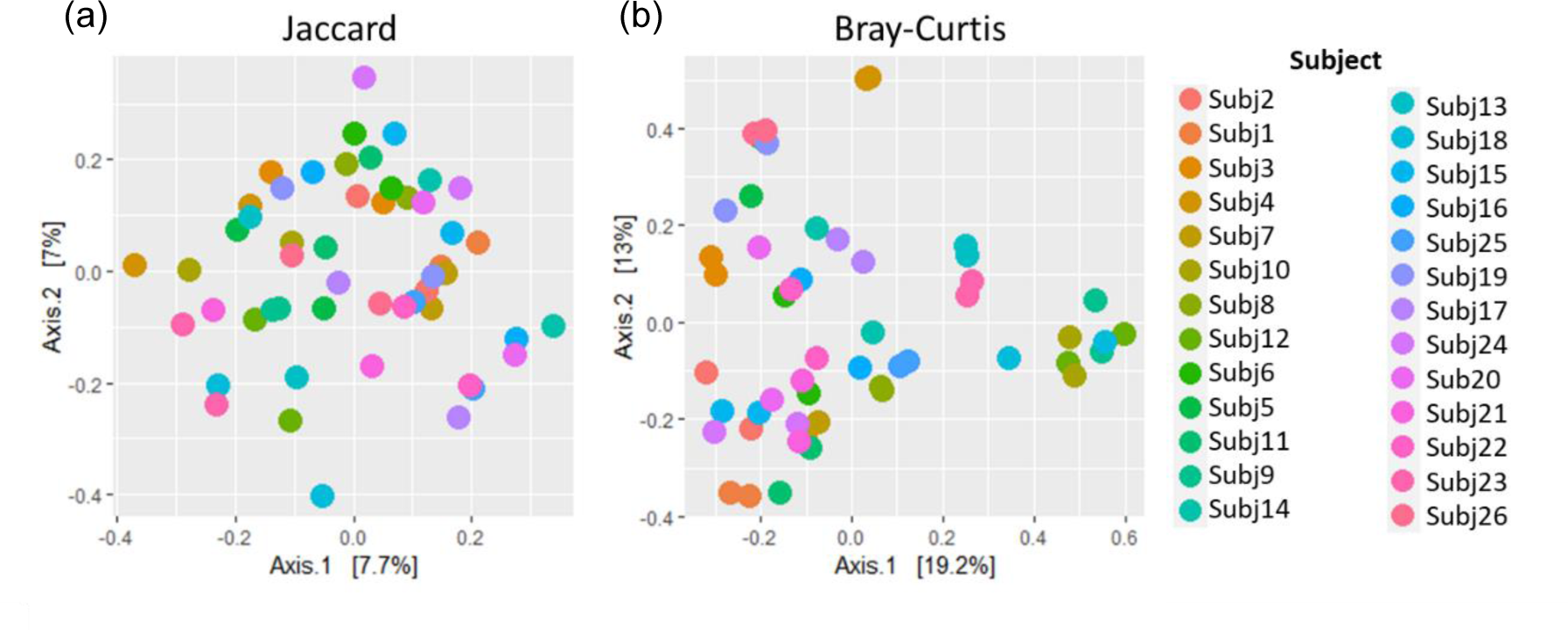
Principal coordinates analysis plots showing (a) Jaccard and (b) Bray Curtis beta diversity measures for pairs of stool and rectal swab samples collected from the same infants at species-level classification.

## 3. Discussion

We have outlined a short-read sequencing protocol which can be used to carry out species-level analysis. Our findings, following analysis of mock controls and technical replicates, indicate that the kits and analytical pipelines used in this study can effectively enable species-level classification and provide reproducible results within and across sequencing plates.

When assessing data from different sample collection approaches using this protocol, our results indicate that care needs to be taken as stool and swab samples collected from the same participant are not comparable at a species level.

### 3.1 Multiple variable regions of 16S rRNA enable adequate species-level analysis

We were able to obtain relatively good species-level resolution in terms of sensitivity using a nearly complete 16S rRNA sequence, which the SNAPP-py3 pipeline developers refer to as a ‘consensus’ sequence. The eight species expected to be found in the ZymoBIOMICS mock controls were not consistently detected to a species level in all mock controls across the seven sequencing plates, suggesting that there is still room for improvement in terms of reproducibility. A previous study comparing the use of different variable regions in Illumina Miseq sequencing, found that using 1-2 variable regions could at best identify 16 of 20 (80%) mock control species correctly (Fouhy et al., 2016). In five mock controls, we detected 7 of 8 mock species (88%), yet for the remaining nine controls we detected all the expected species. Thus, our results would indicate that using all nine variable regions of the 16S rRNA gene improves accuracy.

When focusing only on sequences that were classified to a species level, sensitivity was high in both the mock extraction and mock sequencing controls. However, our results indicate that high sensitivity comes at the expense of obtaining poorer precision in some controls. The overall performance for mock extraction and mock sequencing controls were median F-scores of 0.84 and 0.80 respectively. These results are comparable with other research that obtained F-scores greater than 0.80 using a bioinformatics tool for analysing sequencing data, for which the authors described their results as being indicative of good performance (Özkurt et al., 2022). Poor precision scores in our case were often driven by false positive taxa that were present in very low relative abundances. Thus, removing rare taxa could yield even better results.

The overall percentages of sequences that were classified as expected taxa and to a species level, ranged between 79.11% and 99.98% for mock extraction controls, and between 81.80% and 99.58% for mock sequencing controls. In a study by Szoboszlay et al. (2023) the authors found that between 58.9-68.9% of reads obtained from Illumina sequencing of the V4 region could be correctly classified to a species level. This improved if rare taxa were excluded. Nanopore sequencing of the entire 16S rRNA gene could yield classification of over 81% of reads to a species level (Szoboszlay et al., 2023). Therefore, although we did not remove rare taxa, our protocol could compete with a full-length sequencing approach and perform better than using a single variable region for short-read sequencing. Ultimately, our results indicate that the RDP classifier is able to accurately identify the expected taxa in our controls.

The relative abundances of species in our mock extraction controls sometimes diverged from the theoretical abundances outlined by the suppliers (ZymoBIOMICS), indicating that there may be some bias introduced during the extraction process. O/E ratios provided a quantitative means of assessing the bias introduced in each control. In particular, they assisted with identifying the species for which observed abundances differed most from theoretical abundances and showed that there was greater bias in the mock extraction controls.

There are various stages in 16S rRNA sequencing where prejudice can arise, favouring certain taxa or altering the relative composition of a sample (Nearing et al., 2021). Studies have found that DNA extraction kits and methods can differ in terms of their ability to extract DNA from gram negative and gram positive bacteria, indicating that the results of a study can be influenced depending on which kit and extraction method are used (Videnska et al., 2019, Yuan et al., 2012). The mock sequencing controls (for which already-extracted DNA was obtained from ZymoBIOMICS) more closely resembled the expected abundances. *L. fermentum* differed substantially from the theoretical abundance in the sequencing controls – unlike the extraction controls for which this particular species had followed the expected abundance more closely. This implies that, for the most part, there is minimal bias introduced at the sequencing step. However, there may be some bias specifically in sequencing *L. fermentum*. Our findings complement other studies which have shown that the extraction and amplification of DNA is particularly prone to prejudicing results, while there is less bias introduced at the sequencing stage of microbial research (Brooks et al., 2015, Lee et al., 2012).

Our findings suggest that using this multivariate analysis approach might provide a means to improve the ability of short-read 16S rRNA sequencing studies to study microbial samples at a species level. This approach would benefit from additional ways to minimise bias – particularly in the DNA extraction process.

We considered including a batch effect correction step in the processing of our data. The goal was to reduce bias introduced due to samples being sequenced on different plates and in different runs. However, when visualising the relative abundances of mock sequencing controls, we found that batch effect correction led to far less consistent results across the controls. As our protocol included a plate-wise decontamination step, batch effects would have already been taken into account here. Thus, based on our findings, we considered an additional batch effect correction step excessive and problematic – leading us to exclude this step. Similarly, we found that using a very stringent threshold for decontamination also had a negative impact on our analysis. As such, we would suggest that care needs to be taken when doing additional processing of data, to ensure that bias is not introduced to the data by overcorrecting for contaminants and batch effects.

### 3.2 Short-read multiple variable region analysis can achieve good reproducibility

Observed richness provides an indication of the number of different taxa within samples, while Shannon’s and Simpson’s indices additionally account for how equally represented these taxa are within samples (evenness) (Kers and Saccenti, 2021). Based on our results we have no reason to reject the null hypothesis that there is no difference in alpha diversity measures between within-run technical replicate pairs.

Jaccard’s distance quantifies how diversity varies between samples based on whether or not taxa are present, while Bray Curtis factors in the abundance of taxa (Kers and Saccenti, 2021, Schroeder and Jenkins, 2018). More confidence is usually placed in Bray Curtis compared to Jaccard’s distance (Schroeder and Jenkins, 2018). The similarity between these beta diversity measures for replicate sample pairs was assessed in terms of dICC. dICC is a distance measure which has been established specifically for microbiome data, building on the concept of intraclass correlation coefficients, to look at the similarity between replicate samples (Chen and Zhang, 2022). Higher ICC values are indicative of strong similarity between measures (Koo and Li, 2016), in our case diversity measures for microbial sample pairs. An ICC of 0.5, however, shows moderate reproducibility and a value of < 0.5 suggests poor similarity (Koo and Li, 2016). Thus, our results in which we get dICC values of 0.940 and 0.762 for Bray Curtis and Jaccard respectively, indicate that we get good beta diversity reproducibility when running samples in duplicate on the same plate. Likewise, the diversity of between-run technical replicates was similar with dICC values of 0.899 & 0.597. This suggests that we do not have significant batch effects across sequencing plates of the same run, or across different runs, when following the protocols set out in this study for sequencing and processing short-read multivariate 16S rRNA data. It should be noted that we only had one pair of replicates across runs and this was grouped with the between-plate replicates in what we referred to as our ‘between-run replicates’. Collectively our findings indicate that there is no batch effect introduced. The sequencing and processing of infant stool and rectal swab samples is consistent, providing confidence in the analysis of the microbiome datasets generated using this protocol.

### 3.3 Stool and rectal swab collection approaches are not interchangeable

The implementation of a new kit and pipeline for analysing both stool and rectal swab samples in this study warranted investigation. Stool and rectal swab samples collected from the same infants and at the same time point were found to differ for several diversity measures. Contrary to our findings, studies have suggested that rectal swab samples can be used in place of stool samples and provide a reliable representative measure (Radhakrishnan et al., 2023, Reyman et al., 2019, Bassis et al., 2017). However, a couple of studies exploring diversity and functional roles of the microbiome have previously reported differences between stool and swab samples (Short et al., 2021, Sun et al., 2021). Short et al. (2021) specifically found that beta diversity differed between these sample types, while Sun et al. (2021) found that rectal swab samples varied from stool in terms of both alpha and beta diversity measures. Moreover, there have been indications that differences in the relative abundances of specific taxa may occur between different sample types (Jones et al., 2018). This would be particularly important to consider when comparing the prevalence of individual taxa between groups, such as when using differential abundance analysis.

Furthermore, a study by Bokulich et al. (2019) found that the length of time between the collection and processing of samples is an important factor to take into consideration when using rectal swab samples in sequencing studies. Although samples in their study were ultimately frozen, swabs were posted to the laboratory, while stool samples were immediately placed on ice until they could be collected and taken to the laboratory. Rectal swab samples received and processed after 48 hours following sample collection did not accurately match stool samples – with a higher proportion of *Enterobacteriaceae* being favoured (Bokulich et al., 2019). The stool and rectal swab samples in our study were collected together and stored at -80°C until DNA extraction and sequencing were done, and therefore we do not face the same challenges as Bokulich et al. (2019) in terms of the length of time for which samples remained at room temperature. However, it may be important to consider factors such as the time for which samples are stored in freezers, and the time between DNA extraction and sequencing in future work involving both stool and swab samples.

The fact that swabs were stored in Primestore, while stool was not, would suggest a substantial difference which requires consideration when using a combination of stool and swab samples in carrying out analysis. Several studies have found that bacterial compositions differ across different regions of entire stool samples (Zreloff et al., 2023, Huson et al., 2017, Gorzelak et al., 2015). Spectroscopic analysis has shown that patterns of metabolites can also vary substantially across different regions of stool (Liang et al., 2020, Gratton et al., 2016). Therefore, using only part of the sample does not provide a good characterisation of the entire sample. These studies highlight the importance of homogenising the entire stool sample in order to carry out sequencing (Zreloff et al., 2023, Liang et al., 2020, Gratton et al., 2016). As sequencing of rectal swab samples is not limited to only a component of the collected sample, this sample would be homogenous. This may explain the differences observed between stool and rectal swab sample pairs.

### 3.4 Limitations and future work

The work presented included only one mock extraction and one mock sequencing control on each sequencing plate. Future work could include duplicates of controls on each plate to better assess reproducibility of the controls. It would also expand the options available for statistically comparing these controls.

In this study we evaluated a single pipeline and protocol for analysing data from multiple variable region 16S rRNA sequencing, however in future this protocol would benefit from comparisons to other tools and pipelines. Moreover, different DNA extraction kits and amplicon sequencing kits could be compared to identify the best kits for carrying out multiple variable region analysis. In this analysis we have worked with compositional data, looking at the relative abundances of taxa. However, in future, it would be useful to explore absolute abundances for samples sequenced using this protocol. Finally, we could explore whether other databases – such as databases specific to the human gut microbiome – might enable more effective and accurate classification of consensus sequences from the SNAPP-py3 pipeline.

### 3.5 Conclusion

The results of our 16S rRNA multiple variable region analysis, using short-read Illumina sequencing data from all 9 variable regions, show that there is promise for using this protocol for species-level analysis. We have identified areas for improvement and future work should assess whether this approach is comparable to other bioinformatics pipelines and tools that have been established for analysing multiple variable region short-read data. As we found differences in diversity between stool and swab sample pairs, future analysis of data which has been generated using this protocol should take this into account. Furthermore, our findings indicate that when using new sequencing kits and protocols to study both stool and swab samples, it would be advisable to do analysis to compare pairs of stool and rectal swabs from the same individual to confirm whether these yield comparable results.

## 4. Methods

### 4.1 Extraction controls, sequencing controls and technical replicates

To assess the reproducibility and accuracy of DNA extraction and sequencing steps, mock controls and technical replicates were included on each plate (Figure 9). A ZymoBIOMICS™ Microbial Community Standard (catalog number ZR D6300), consisting of eight known bacterial species including both gram-positive and gram-negative bacteria, was included on each plate as a DNA extraction control. Each plate included a ZymoBIOMICS™ Microbial Community DNA Standard (catalog number ZR D6305), which contains already extracted DNA for eight known bacterial species. This served as a sequencing control. Seventeen within-run and 8 between-run technical replicate pairs were also included across the sequencing plates.

**Figure 9:**
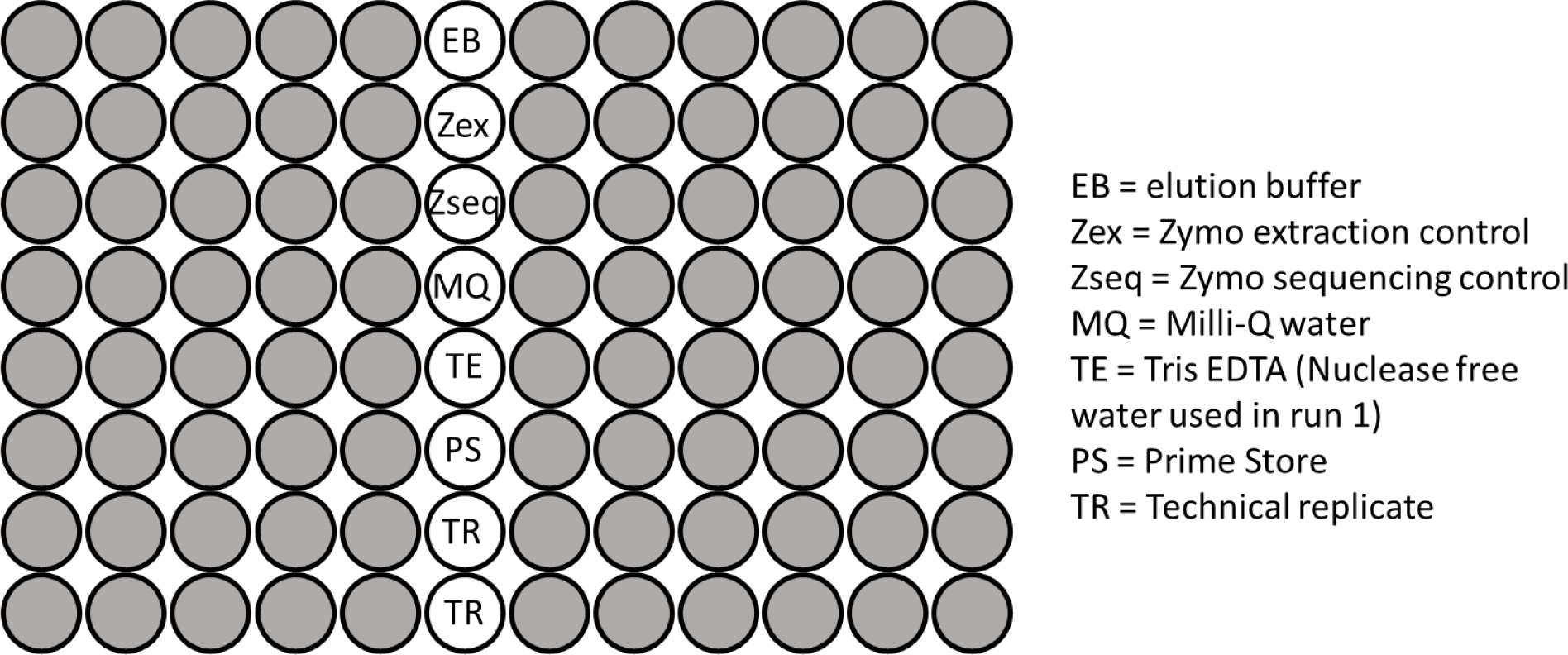
The plate layout used for sequencing microbial samples, including the negative controls, positive (mock) controls and replicates included.

### 4.2 Stool and swab sample collection

Twenty six pairs of stool and rectal swab samples were collected at the age of 0-5 weeks, for infants from the Healthy Baby Study born between 37 and 42 weeks gestational age. Swabs were stored in Primestore solution (PrimeStore® Molecular Transport medium). All stool and swab samples were transferred to a -80°C freezer for long-term storage.

Stool samples were thawed, and half of a pea-sized scoop was collected from the side/centre of the sample. This was placed in a tube with 750µl of lysis buffer. For rectal swab samples, 400µl of sample in Primestore was placed in a tube with 400µl of lysis buffer. These then underwent off-board lysis, using the QT Qiagen bead beater, prior to DNA extraction.

### 4.3 DNA extraction, preparation of sequencing library and Illumina sequencing

Manual DNA extraction of the stool and rectal swab samples and mock extraction controls was carried out using the Quick-DNA^TM^ Fecal/Soil Microbe Microprep Kit (ZymoBIOMICS catalog number D6012). Prior to carrying out polymerase chain reaction (PCR), Qubit^TM^ (Thermo Fisher Scientific, Waltham, MA, USA) was done to check the starting DNA concentrations. xGen^TM^ 16S Amplicon Panel v2 kits (Integrated DNA Technologies, Coralville, IA, USA) were used in library preparation for sequencing. Kits included primer pairs for amplification of all nine hypervariable regions of the 16S rRNA gene. Additionally, the primers for these kits have dual indices to allow greater numbers of samples to be run together in a single flow cell. Moreover, the xGen kits include Normalase^TM^ which could be used to enzymatically normalise library sizes prior to sequencing. Finally, qPCR was performed following the Normalase step to quantify the final library size prior to sequencing.

Negative controls, including Primestore, Milli-Q water, elution buffer and Tris EDTA/Nuclease free water, were added to each plate (Figure 9) together with prepared libraries from the stool and rectal swab samples. Mock controls and technical replicates, as described above, were also included on each plate. Sequencing was conducted across two sequencing runs on seven plates using an Illumina MiSeq platform (Illumina, San Diego, CA, USA). The combined 16S library per run was subjected to paired-end sequencing on the Illumina® MiSeq™ platform, employing the MiSeq Reagent v3 kit with 600 cycles (Illumina, San Diego, CA, USA).

### 4.4 Bioinformatics processing of sequencing data

Ethics approval for analysis of this data was provided by the Human Research Ethics Committee, University of Cape Town (557/2020). Following sequencing, preprocessing steps were carried out for quality control and to prepare the data for statistical analysis (Figure 10). Raw sequencing data was run through FastQC (Andrews, 2010) to assess the quality of the reads. Following this quality control step, forward and reverse reads for each sample were processed using the SNAPP-py3 pipeline (Chai, 2021). Sequencing data from run 1 and run 2 were processed separately. Of the four main output files from the pipeline, the lineage table and an adapted taxonomy table were used for further analysis.

**Figure 10:**
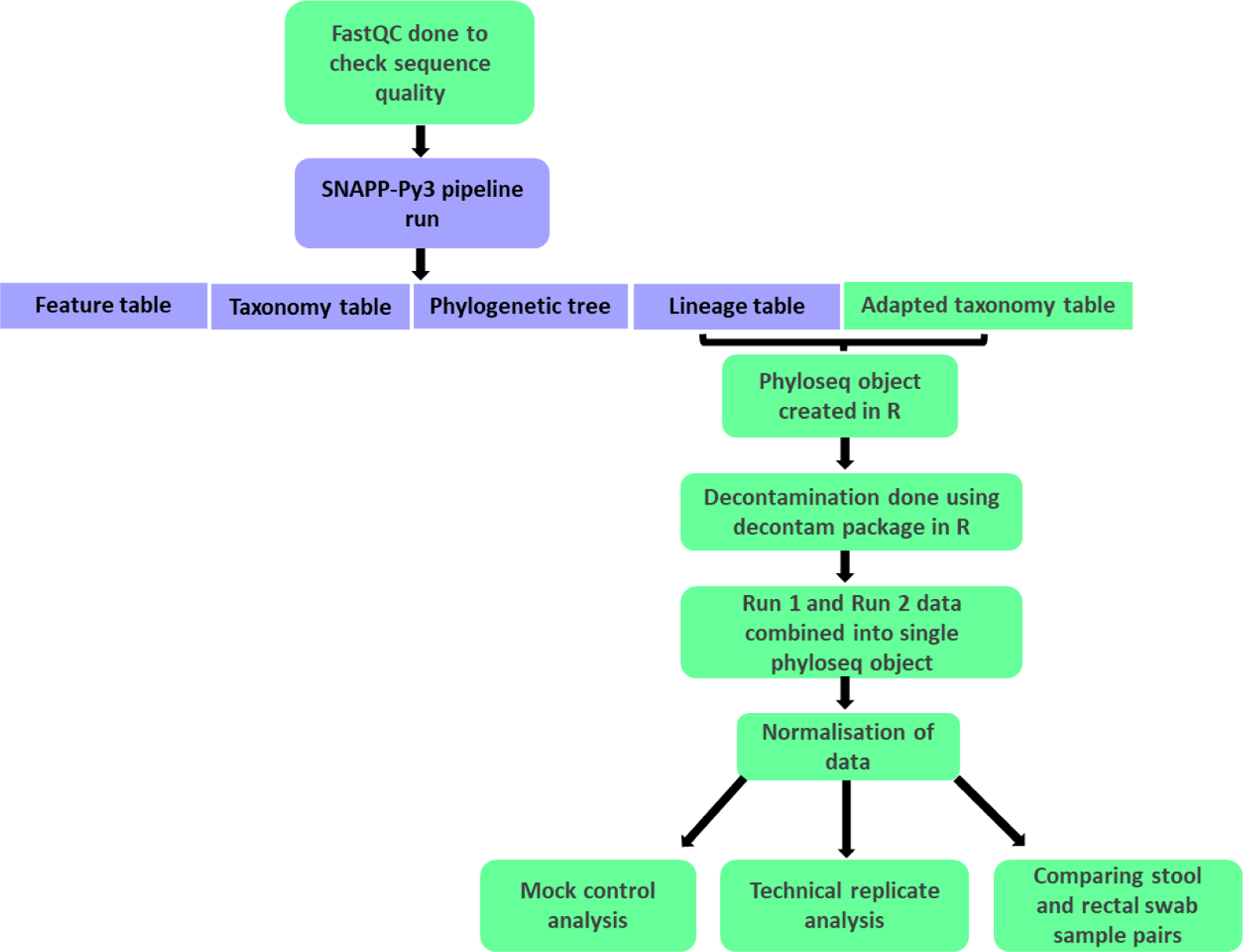
A flowchart summarising the processing and analysis steps carried out using raw forward and reverse read sequencing files. Boxes in green indicate the steps included in addition to the main SNAPP-py3 pipeline.

The remaining processing and analysis were carried out in R version 4.2.1 (R Core Team, 2022). This stage of processing began with creating a phyloseq object (McMurdie and Holmes, 2015, McMurdie and Holmes, 2013). Using the decontam package (Davis et al., 2018), decontamination was carried out separately for each plate, using plate-specific negative controls. A combined frequency and prevalence approach was used, selecting a threshold of 0.1 for the prevalence component. Phyloseq objects from runs 1 and 2 were then combined into a single phyloseq object for further downstream processing and analysis.

A normalisation step to account for different library sizes was implemented by determining the median library size and normalising each sample accordingly (Balle et al., 2020, The Jackson Laboratory, 2019). Finally, subsetting into various phyloseq objects was done to prepare for downstream statistical analysis. We ultimately had separate phyloseq objects containing the mock extraction controls, mock sequencing controls, within-run repeats, between-run repeats and baseline pairs of stool and rectal swab samples.

Batch effect correction was done using MMUPHin (Ma, 2022, Ma et al., 2022). This data was compared to data in which no batch effect correction was carried out, to determine the necessity of accounting for batch effects.

### 4.5 Statistical analysis

Relative abundances for controls, replicates and stool/rectal swab sample pairs were visualised using QIIME2 software (Bolyen et al., 2019). All microbiome analysis was done at a species level in R. Genus-level and phylum-level analyses were additionally included in select steps to provide additional insights.

Performance measures were calculated as outlined by Özkurt et al. (2022) using data for sequences classified to a species-level. Precision and sensitivity were calculated based on the number of correctly identified species expected to be in the mock control (true positives [TP]), the number of expected species that were not detected (false negatives) and the number of non-expected species classified as being in the control (false positive [fp]). F-scores could then be calculated based on these values. The calculations used were as follows:

Precision = TP/(TP+FP)

Sensitivity = TP/(TP+FN)

F-score = 2*precision*sensitivity/(precision+sensitivity) (Özkurt et al., 2022).

The percentage relative abundances of each of the eight expected species were determined for mock cell (extraction) and mock DNA (sequencing) controls on each sequencing plate. For each control, the total percentage of sequences that were correctly classified as an expected species, was calculated. Furthermore, observed relative abundances were compared to the theoretical abundances provided by ZymoBIOMICS for each species in the mock controls by calculating Observed/Expected (O/E) ratios (Maki et al., 2023).

Functions from the phyloseq package in R (McMurdie and Holmes, 2015, McMurdie and Holmes, 2013), specifically the estimate_distance and distance function, were used to calculate alpha and beta diversity measures for replicates and stool/rectal swab samples. The alpha diversity measures included are observed richness (Fisher et al., 1943), Shannon’s index (Shannon, 1948) and Simpson’s index (Simpson, 1949). Bray Curtis (Bray and Curtis, 1957) and Jaccard’s (Ludwig and Reynolds, 1988) distances were the beta diversity measures included in our analysis. Plot_richness and plot_ordination functions were used to plot alpha and beta diversity measures, respectively. Paired Wilcoxon tests were used to compare alpha diversity measures between pairs of within-run replicates and to identify differences between pairs of between-run replicates. In order to determine whether beta diversity measures were reproducible between technical replicate pairs, distance-based intraclass correlation coefficients (dICCs) were calculated separately for within-run and between-run replicates (Chen and Zhang, 2022). Paired Wilcoxon tests and dICCs were similarly used to compare pairs of stool and rectal swab samples collected from the same infants.

## Supporting information

Code for supporting data

## Acknowledgements

We would like to acknowledge and thank Dr Veronica Allen for assisting Dr Fadheela Patel with DNA extraction and the 16S rRNA sequencing. We thank Dr Samantha Fry and Thandiwe Hamana for organizing visits and the sometimes difficult task of sample collection. We thank Dr Farai Mberi and Caylin Mc Farlane for their assistance with sample transportation and storage. Our sincere thanks to Slindile Mbhele for all she did to arrange the storage and transportation of samples as project manager. Thank you also to Dr Benli Chai, the developer of the SNAPP-py3 pipeline, for providing assistance and advice.

## Data availability

Data has been prepared in excel spreadsheets which will be made available with this manuscript. Unique identifying numbers have been replaced with new sequentially assigned identifiers for this manuscript.

## Author contributions

AG carried out the analysis and drafted this manuscript. MH, MK and AvdK, as Principal Investigators, designed and oversaw the larger microbiome study. FP carried out the sequencing of the controls, replicates and samples. FL provided guidance with the statistical analysis. All co-authors assisted with editing the manuscript.

