## Supplementary figures & tables for "Using short-read 16S rRNA sequencing of multiple variable regions to generate high-quality results to a species level"

**Supplementary material**

**Supplementary table 1:** The expected theoretical abundances and relative abundances of the eight expected bacterial genera in mock cell controls given as percentages. The total percentage of correctly classified sequences for each control is also given.

| Genera | Theoretical abundance (%) | R1P1  Zymoex | R1P2  Zymoex | R1P3  Zymoex | R1P4  Zymoex | R2P1  Zymoex | R2P2  Zymoex | R2P3  Zymoex |
| --- | --- | --- | --- | --- | --- | --- | --- | --- |
| Limosilactobacillus | 18.4 | 18.21 | 18.27 | 17.99 | 17.64 | 18.61 | 16.60 | 18.02 |
| Bacillus | 17.4 | 18.40 | 19.55 | 19.72 | 20.62 | 18.77 | 17.63 | 18.55 |
| Staphylococcus | 15.5 | 10.84 | 9.17 | 9.14 | 11.50 | 10.97 | 9.32 | 10.76 |
| Listeria | 14.1 | 4.42 | 4.47 | 4.59 | 4.97 | 0.00 | 4.59 | 4.67 |
| Salmonella | 10.4 | 17.58 | 18.90 | 17.61 | 16.37 | 10.95 | 20.39 | 19.22 |
| Escherichia/Shigella | 10.1 | 17.08 | 18.31 | 18.26 | 18.91 | 20.58 | 19.26 | 20.50 |
| Enterococcus | 9.9 | 5.39 | 4.53 | 4.59 | 5.28 | 5.36 | 4.96 | 4.81 |
| Pseudomonas | 4.2 | 7.37 | 6.53 | 7.80 | 4.51 | 4.63 | 7.25 | 3.40 |
| Other | 0 | 0.71 | 0.27 | 0.31 | 0.21 | 10.14 | 0.01 | 0.07 |
| % Correctly classified |  | 99.29 | 99.73 | 99.69 | 99.79 | 89.86 | 99.99 | 99.93 |

**Supplementary table 2:** The expected theoretical abundances and relative abundances of the eight expected bacterial genera in mock DNA controls given as percentages. The total percentage of correctly classified sequences for each control is also given.

| Genera | Theoretical abundance (%) | R1P1  Zymoseq | R1P2  Zymoseq | R1P3  Zymoseq | R1P4  Zymoseq | R2P1  Zymoseq | R2P2  Zymoseq | R2P3  Zymoseq |
| --- | --- | --- | --- | --- | --- | --- | --- | --- |
| Limosilactobacillus | 18.4 | 13.08 | 12.60 | 13.36 | 11.97 | 13.89 | 13.16 | 12.86 |
| Bacillus | 17.4 | 17.89 | 17.85 | 17.44 | 17.05 | 17.81 | 17.66 | 17.99 |
| Staphylococcus | 15.5 | 13.85 | 13.42 | 13.91 | 14.11 | 13.96 | 14.22 | 14.04 |
| Listeria | 14.1 | 15.28 | 14.81 | 14.60 | 15.43 | 15.85 | 15.95 | 16.06 |
| Salmonella | 10.4 | 11.49 | 12.04 | 13.12 | 6.32 | 0.01 | 11.23 | 9.06 |
| Escherichia/Shigella | 10.1 | 11.59 | 12.40 | 10.86 | 11.63 | 12.18 | 11.20 | 11.65 |
| Enterococcus | 9.9 | 11.98 | 11.18 | 11.54 | 11.16 | 11.89 | 12.40 | 11.76 |
| Pseudomonas | 4.2 | 4.45 | 5.11 | 4.66 | 4.98 | 3.72 | 4.09 | 3.41 |
| Other | 0 | 0.39 | 0.59 | 0.51 | 7.35 | 10.70 | 0.08 | 3.17 |
| % Correctly classified |  | 99.61 | 99.41 | 99.49 | 92.65 | 89.30 | 99.92 | 96.83 |

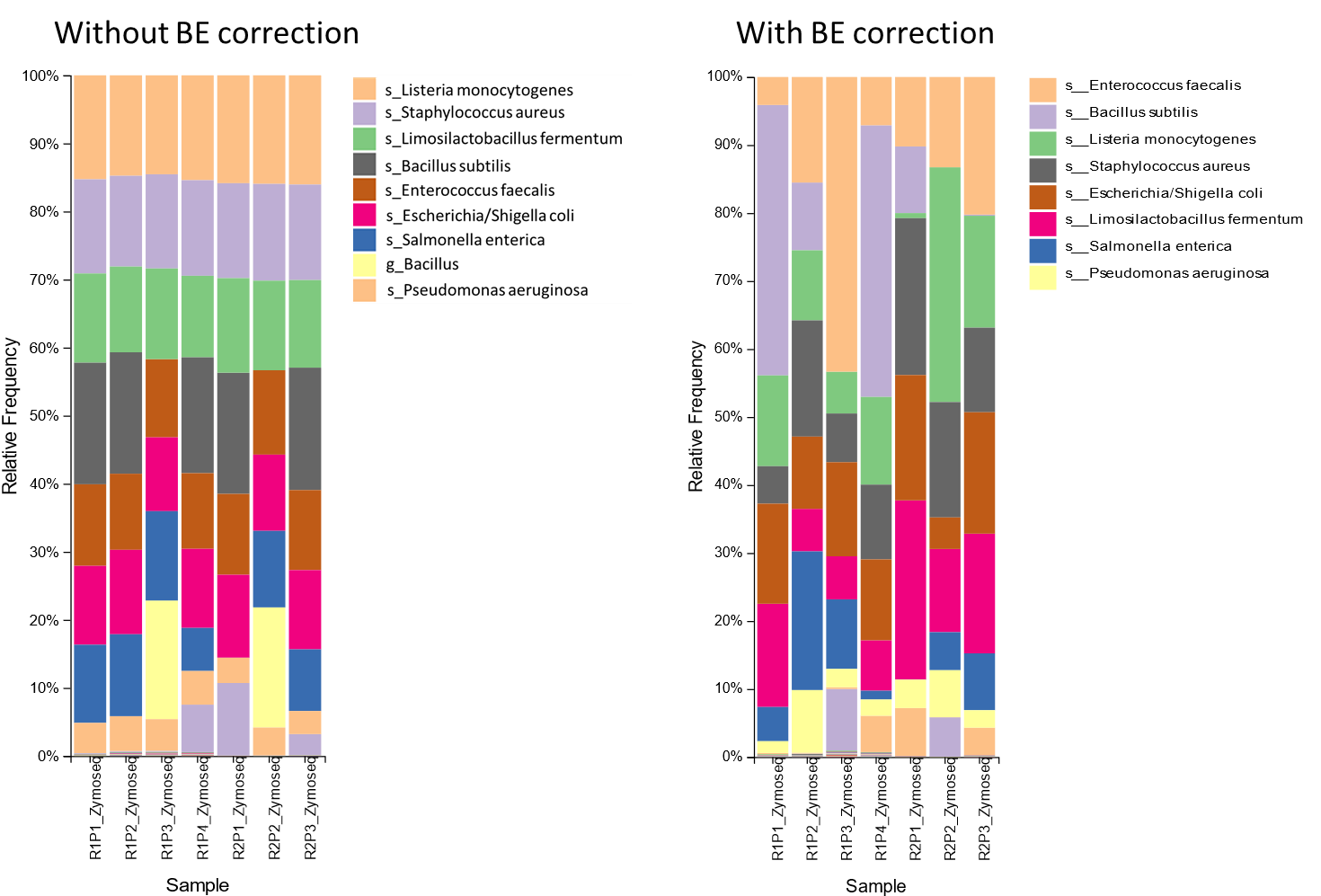

(a)

(b)

**Supplementary figure 1**: Relative abundances of mock sequencing controls (a) before and (b) after batch effect correction carried out with MMUPHin. Legend: s = species, g = genus level classification.

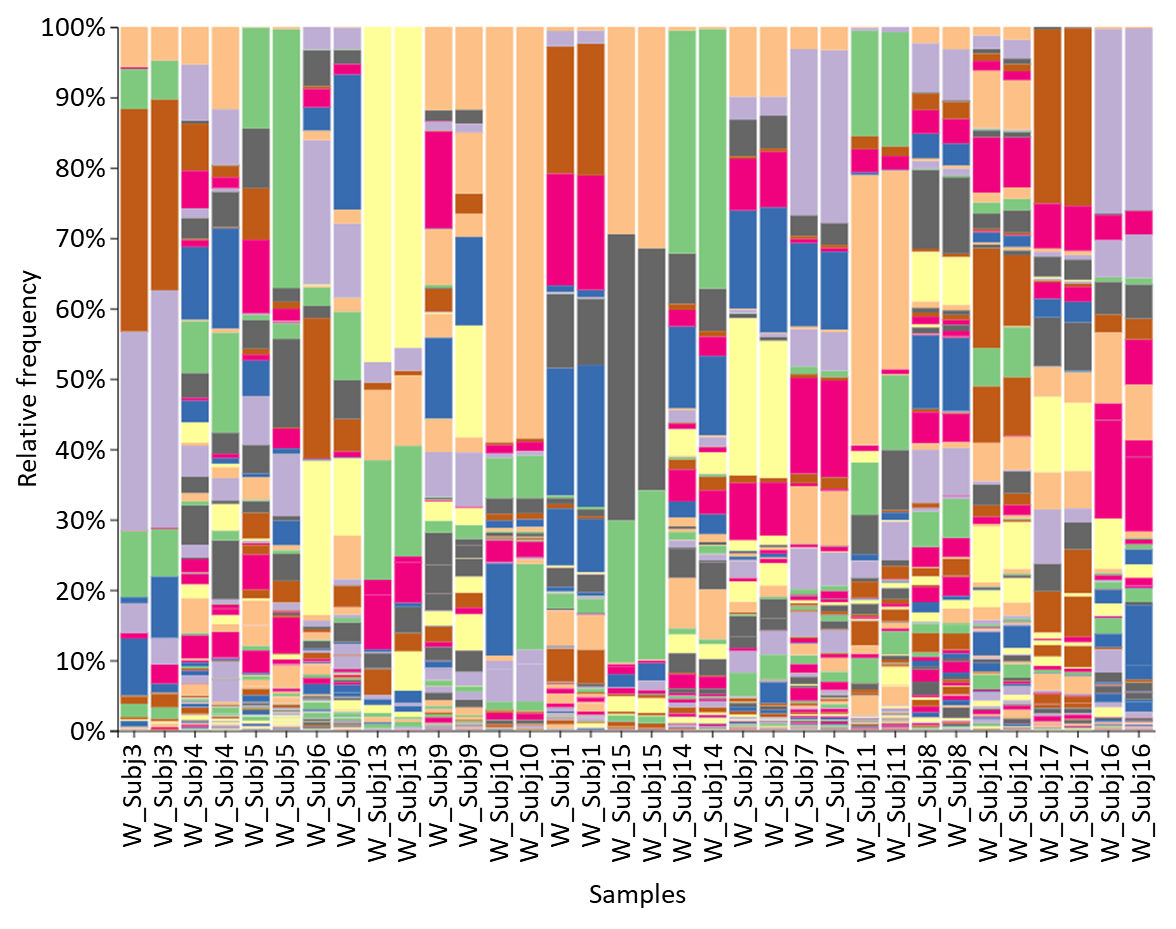

**Supplementary figure 2:** Relative abundance of species for within-run repeats.

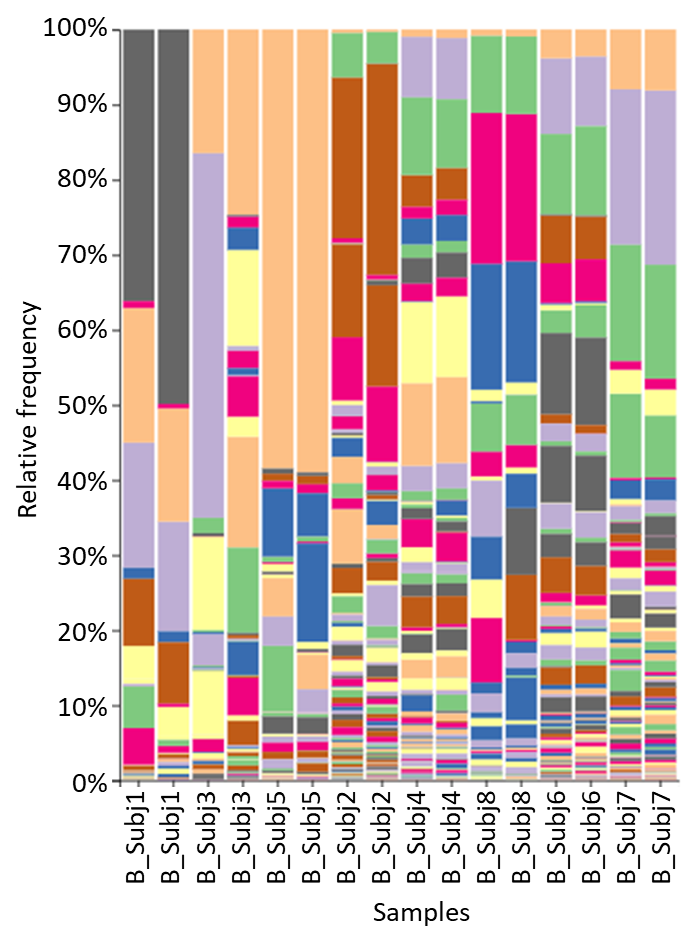

**Supplementary figure 3:** Relative abundance of species for between-run repeats

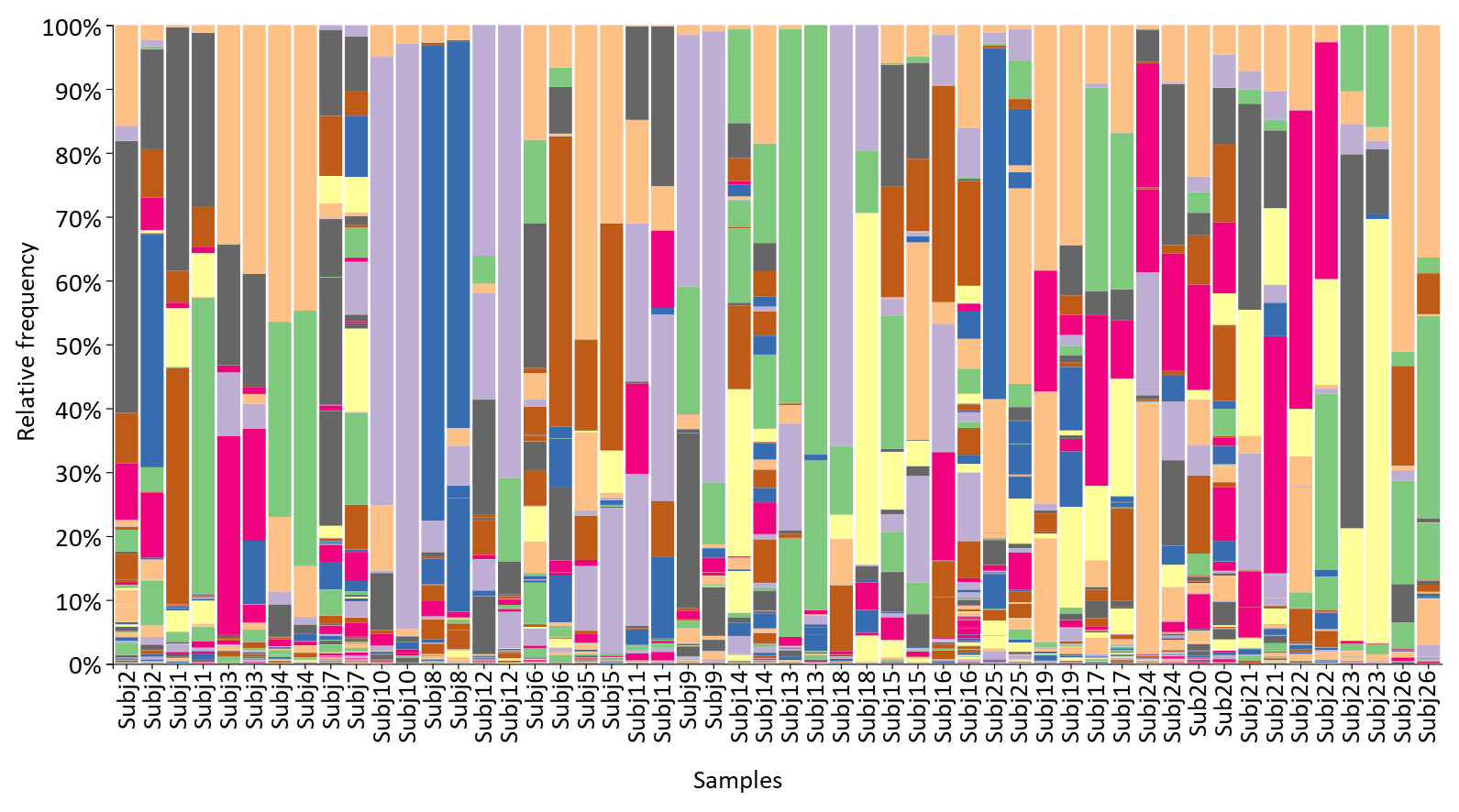

**Supplementary figure 4:** Relative abundance of species for baseline stool and swab sample pairs collected from the same infants.
