## Supplementary material for "Using short-read 16S rRNA sequencing of multiple variable regions to generate high-quality results to a species level": Code for supporting data

**Creating a phyloseq object using the supporting data excel spreadsheets**

In order to create a phyloseq object in R, 3 different sets of data are needed: a matrix of counts, taxonomy information and metadata. Therefore, there are 15 supporting data spreadsheets - 3 each for the mock extraction, mock sequencing, within-run repeats, between-run repeats and stool/swab analysis.

#Showing an example for reading the mock DNA control spreadsheets into R and creating a phyloseq object for downstream analysis.

#Reading in the excel spreadsheets for mock controls

mock_dna_count <- read.xlsx(file="mock_dna_count.xlsx", header=TRUE, sheetName="sheet1", row.names = 1)

mock_dna_tax <- read.xlsx(file="mock_dna_tax.xlsx", header=TRUE, sheetName="sheet1", row.names = 1)

mock_dna_samples <- read.xlsx(file="mock_dna_samples.xlsx", header=TRUE, sheetName="sheet1", row.names = 1)

table(rownames(mock_dna_tax) == rownames(mock_dna_count))

mock_dna_tax_matrix <- as.matrix(mock_dna_tax)

mock_dna_count_matrix <- as.matrix(mock_dna_count)

#Creating a phyloseq object

library(phyloseq)

phy <- phyloseq(otu_table(mock_dna_count_matrix, taxa_are_rows = T),tax_table(mock_dna_tax_matrix), sample_data(mock_dna_samples))

str(phy)

phy
